# Tipping-point analysis uncovers critical transition signals from gene expression profiles

**DOI:** 10.1101/668442

**Authors:** Xinan H Yang, Zhezhen Wang, Andrew Goldstein, Yuxi Sun, Megan Rowton, Yanqiu Wang, Dannie Griggs, Ivan Moskowitz, John M Cunningham

## Abstract

Differentiation involves bifurcations between discrete cell states, each defined by a distinct gene expression profile. Single-cell RNA profiling allows the detection of bifurcations. However, while current methods capture these events, they do not identify characteristic gene signals. Here we show that BioTIP – a tipping-point theory-based analysis – can accurately, robustly, and reliably identify critical transition signals (CTSs). A CTS is a small group of genes with high covariance in expression that mark the cells approaching a bifurcation. We validated its accuracy in the cardiogenesis with known a tipping point and demonstrated the identified CTSs contain verified differentiation-driving transcription factors. We then demonstrated the application on a published mouse gastrulation dataset, validated the predicted CTSs using independent in-vivo samples, and inferred the key developing mesoderm regulator Etv2. Taken together, BioTIP is broadly applicable for the characterization of the plasticity, heterogeneity, and rapid switches in developmental processes, particularly in single-cell data analysis.

**Highlights:**

- Identifying significant critical transition signals (CTSs) from expression noise
- A significant CTS contains or is targeted by key transcription factors
- BioTIP identifies CTSs accurately and independent of trajectory topologies
- Significant CTSs reproducibly indicate bifurcations across datasets

## 1. Introduction

Fate decisions in stem cell differentiation involve bifurcations between discrete cellular states that present distinct gene expression profiles. Before a bifurcation, multilineage progenitor cells undergo priming, the phenomenon of cells in a subpopulation that are permissive for opposing cell fates prior to their lineage commitment (Moris et al., 2016; Ranzoni et al., 2021; Teschendorff and Feinberg, 2021). Besides epigenetic priming, transcriptional priming has been recently described using single-cell transcriptomes (Ando et al., 2019; Mojtahedi et al., 2016; Nestorowa et al., 2016; Zhao and Choi, 2019; Zhou et al., 2019). Transcriptional priming are detectable prior to two types of bifurcations (Teschendorff and Feinberg, 2021) (**Fig 1a**). Pitchfork bifurcations happen before a multipotential progenitor cell makes a quick bifurcation between two distinct stable states (Bargaje et al., 2017); saddle-node bifurcations occur before a progenitor cell approaches a threshold of irreversible commitment to a differentiated state (Mojtahedi et al., 2016; Richard et al., 2016). However, mainstream analyses assumed that pluripotent cells undergo smooth and continuous transitions to committed states that are marked by the progressive activation and silencing of molecular hallmarks (Lummertz da Rocha et al., 2018), being unable to characterize these abrupt, non-linear, and often irreversible transition events. Therefore, it is crucial to fathom where bifurcations lie and, importantly, what characterizes or drives their formation.

**Figure 1.**
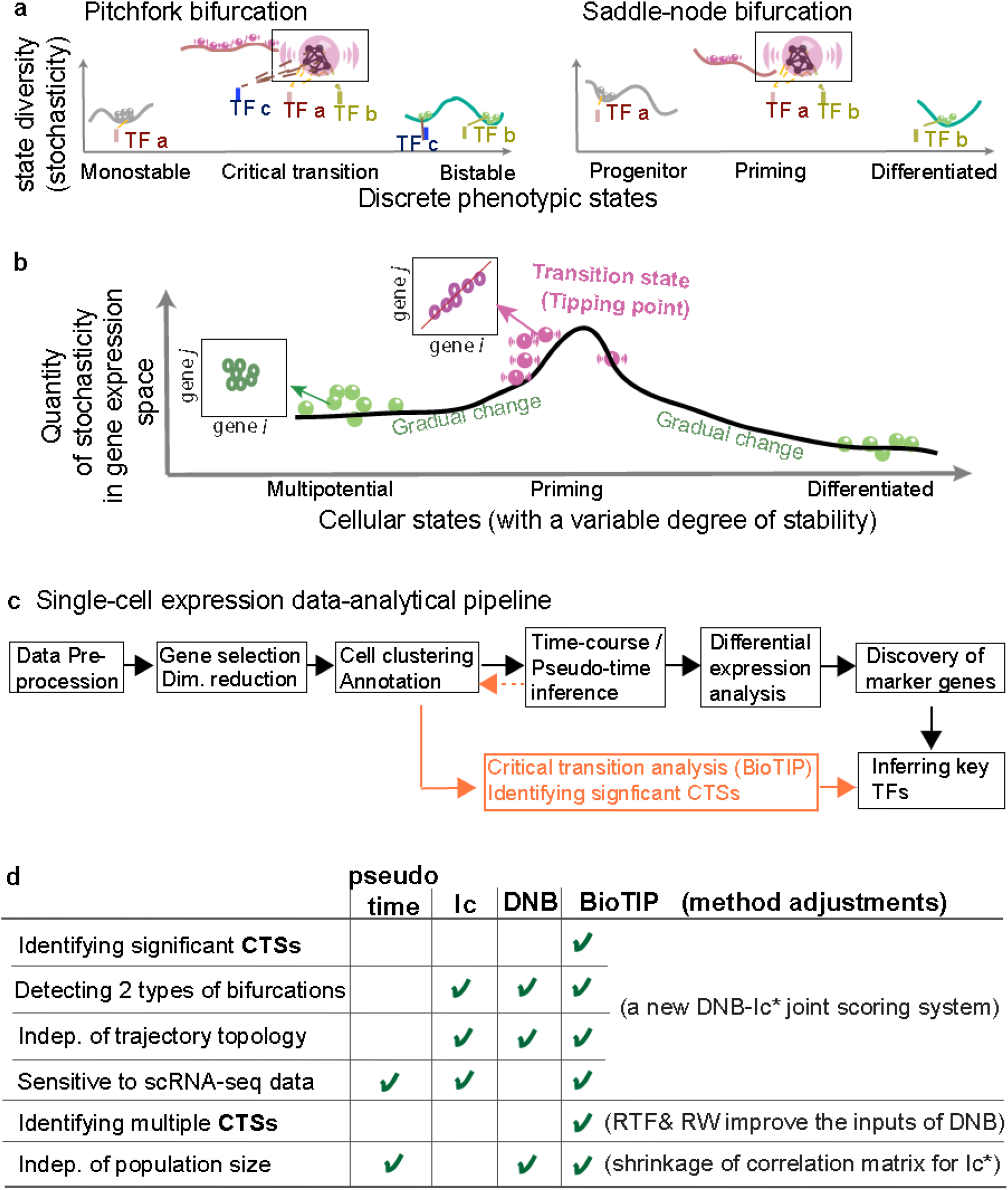
Tipping-point theory and adaption to gene expression analysis. **a**, A model of transcription factor (TF)-regulated stochasticity described in two formats: the pitchfork bifurcation (e.g., lineage bifurcation, left) and the saddle-node bifurcation (e.g., irreverent fate commitment, right). In this model, discrete cell states present distinct gene expression patterns, each under the control of particular TFs. According to tipping-point theory, a fixed point (a basin with steep local gradient near the center) is illustrated with a line, and the systems (cells) are illustrated with balls. At the critical transition (tipping point), the line turns flat, representing the effect of multiple decayed attractions resultant in a vulnerable regulation. A transcriptional readout of this phenomena is the increased covariance among a small set of interacting genes (the circles in the embedded box). **b**, Schematic of the gene expression space of a system undergoing a bifurcation, exampled by a transition from multipotential progenitor state to a differentiated stable state. A tipping point in this progression occurs when simultaneous emergence of new controlling transcription factors increases the diversity/stochasticity in a cell population (middle panel). Before making a cell fate decision, an onset of correlated changes in expression among a small set of genes precipitate the disequilibrium (red shaking balls), which is illustrated by the pink cycles in the embedded box. After passing this tipping point, system diversity drops quickly and remains lower compared with the original progenitor state. **c**, Analytic pipeline of single-cell RNA-seq analysis, with BioTIP as an alternative to differential expression analysis. Arrow lines in orange showing that BioTIP is applied to cell state ensembles (clusters) without the pseudo-order information. The dashed orange line showing that BioTIP may use pseudo-order information to select cell states on the trajectory of interests. **d**, Table view comparing the functionality of BioTIP with pseudo-time analysis and two existing tipping-point scoring systems. DNB: Dynamic biomarker network; Ic: index of criticality; Ic*: a redefined Ic; RTF: relative transcript fluctuation; RW: random walk.

To detect where bifurcations lie using gene expression profiles (Mojtahedi et al., 2016; Richard et al., 2016), researchers have adopted a theory called ‘tipping point’ that has wide applications in ecosystems, climates, and other complex systems (Clements et al., 2019; Lenton and Livina, 2016). Bifurcations (a type of tipping point) occur when perturbations in some gene interactions causes the cell to cross a certain threshold that had been maintaining an equilibrium (Teschendorff and Feinberg, 2021). According to this theory, when a critical transition is imminent, a set of interconnected genes gain covariance (due to the loss of the preexisting equilibrium and the onset of a new state) (**Fig 1b**, embedded red dots) (Chen et al., 2012; Clements et al., 2019; Teschendorff and Feinberg, 2021). Knowing this, the expression patterns of these genes thus serve as a critical transition signal (**CTS)** to characterize an impending bifurcation.

There are two models about what drives the formation of critical transitions: ‘regulated stochasticity’ (Teschendorff and Feinberg, 2021) and stochasticity. The regulated-stochasticity model assumes that extrinsic inputs alter the expression of CTS genes to change cell states. Established bifurcation-driving inputs include transcription factor (**TF**) whose temporal activities determinate distinct cell fates (Bockamp et al., 1998; Hu et al., 1997; Shai et al., 2015; Sheikh and Groom, 2020), which is illustrated in **Figure 1a**. The stochasticity model assumes that stochastic fluctuations in a single cell (which is hereafter called **transcriptional noise**) cause cells to switch between states. The stochasticity model is supported by the transitions displaying stochastic fluctuations, e.g., cell’s position in the cell cycle phases (Antolovic et al., 2019; Buganim et al., 2012; Kalmar et al., 2009). In the extreme, the second model anticipates overall gene expression changes to participate in the state transition, i.e., the model does not identify CTSs.

To identify CTSs, current computational applications have uniformly been applied to bulk RNA-seq data (Chen et al., 2012; Richard et al., 2016; Yang et al., 2015). However, bulk samples pose challenges for the significance and reproducibility in the TF-regulated CTS analysis. This is because transcriptional regulation fluctuates spatiotemporally in living cells, but the heterogeneity of single cells is erased in bulk samples. By addressing the challenges arising for single-cell RNA sequencing (**scRNA-seq**) (**Fig 1c**), we aim to identify significant and robust CTSs.

We introduce **BioTIP**, a new method that complements mainstream scRNA-seq analytical methods (Delile et al., 2019; Trapnell et al., 2014) (**Fig 1c**). BioTIP’s performance was evaluated through application to both simulated and benchmark datasets and compared to previous analyses. We demonstrated that not only can BioTIP be applied to the characterizations of bifurcations, but also to address the computational challenges in analyzing single-cell transcriptomes. BioTIP was then applied to the scRNA-seq data of mouse gastrulation on which tipping point analyses had never been done before, where multiple critical transitions were identified and evaluated using independent cells of the same developmental stages. Based on literature, we showed how a significant CTS provides insight into cell-fate decisive transcription factors. These results open a way for characterizing critical transitions from dynamic expression profiles of developmental biology.

## 2. Results

### 2.1. BioTIP overview

Temporally transcriptional regulation must be understood at the level of the single cell (Richard et al., 2016). However, classical tipping-point models require a time series in which the same cell is analyzed at various successive points in time (Clements et al., 2019; Lenton and Livina, 2016). This type of longitudinal transcriptomic data is rarely available for single-cell transcriptomes. In the absence of longitudinal data, a common practice has been to describe each cellular state from its statistical replicates (a cluster of individual cells sharing similar gene expression patterns) (Chen et al., 2012; Mojtahedi et al., 2016). In these practices, the cell clusters composed of bifurcation-occurring or impending cells were defined as ‘critical transition states’ or ‘tipping points.’ Therefore, we hereafter rename the classical ‘early-warning signal for tipping point’ (which suggests time series) as a ‘<u>c</u>ritical-<u>t</u>ransition <u>s</u>ignal’ (**CTS**). The term CTS encourages the application of tipping-point theory with broadly available scRNA-seq profiles.

To adopt tipping-point theory to single-cell transcriptomic analysis, two assumptions must be met:

1. The system (an ensemble of the individual replicates) has a dissipative structure (i.e., having discrete states including the one showing semi-stability).
2. Each state has a characteristic gene expression profile and thus presents a distinct molecular phenotype.

Complementing to the current scRNA-seq analytical pipeline, BioTIP focuses on gene expression dynamics and is applicable to cellular states regardless of their pseudo-orders (**Fig 1c**). For scRNA-seq data, BioTIP is unique in its ability to address the five following challenges in the identification of significant CTS (**Fig 1d**):

a. Bifurcations could be stimulated by signals other than TFs, particularly when scores of random genes indicate bifurcations (Antolovic et al., 2019; Buganim et al., 2012; Kalmar et al., 2009), thus noise removal is essential.
b. A significance threshold is required because increased gene expression stochasticity could indicate tipping points (Richard et al., 2016), and multiple CTSs and bifurcations may coexist.
c. Because we can profile only subsets of the true trajectories, the independence of trajectory topologies is important, particularly for identifying saddle-node bifurcations.
d. Due to the high resolution of cell states detectable, changes of certain genes could span several states of distinct phenotypes and gene expression patterns.
e. The sizes of the statistical ensembles (e.g., cellular populations) vary considerably.

To identify CTS from noisy gene expression profiles, the regulated-stochasticity model-based Dynamic Network Biomarker (**DNB**) is the most promising and first method to have come into use (Chen et al., 2012; Liu et al., 2017; Yang et al., 2018; Yang et al., 2015; Zhang et al., 2019). DNB predicts a tipping point from which the CTS is detected. Therefore, DNB enables transcriptomic analysis on sample ensembles of macroscopic phenotypes, allowing for both time-series and cross-sectional analyses with one model. However, DNB may report false positive from complex scRNA-seq data because the method compares gene modules at each state over all states (**Methods, Formula 4**). This comparison is susceptive to genes that covariate in multiple states and can’t identify multiple CTSs. To overcome these limits, BioTIP designed two enhanced gene-selections steps to better screen inputs for DNB. Additionally, BioTIP inputs the DNB-reported CTS candidates into a stochasticity model-based scoring system for the significance evaluation. The rational comes from the observed agreement between these two models in one cell system (Richard et al., 2016).

Among existing stochasticity model-based methods, index of criticality (**Ic**) is a breakthrough in detecting bifurcations using single-cell transcriptomes (Mojtahedi et al., 2016). However, Ic is prone to overpredict states of small size (i.e., with small number of replicates) as critical transitions. This is a result of an inaccurate numerator in the Ic ratio which inherits defects from the classical estimation of correlation matrix that inflates for states with small sample sizes (See Methods). To overcome this, we redefine an Ic* score with a ‘shrinkage of correlation’ method (Schafer and Strimmer, 2005).

Altogether, BioTIP analysis has five steps that are grouped into three main components (**Fig 2a, Methods**). The first component (step i) searches for critical transitions from noisy background signals using the new Ic* scoring system (**Methods, Formulas 1-2**). Being served with a newly designed score (**Formula 3**) to select variable genes robustly and to uncover co-expressed genes fast, the second component (steps ii-iv) detects CTS candidates based on the existing DNB method (**Formula 4**). The final component (step v) uses a Delta score to identify significant CTSs from candidates. The Delta score is based on where the maximum Ic* occurs and its distance from the second maximum Ic* (**Formula 5**). Together, we can assess the significance of each CTS candidate meeting following three quantitative criteria, enabling the identification of multiple CTSs.

**Figure 2.**
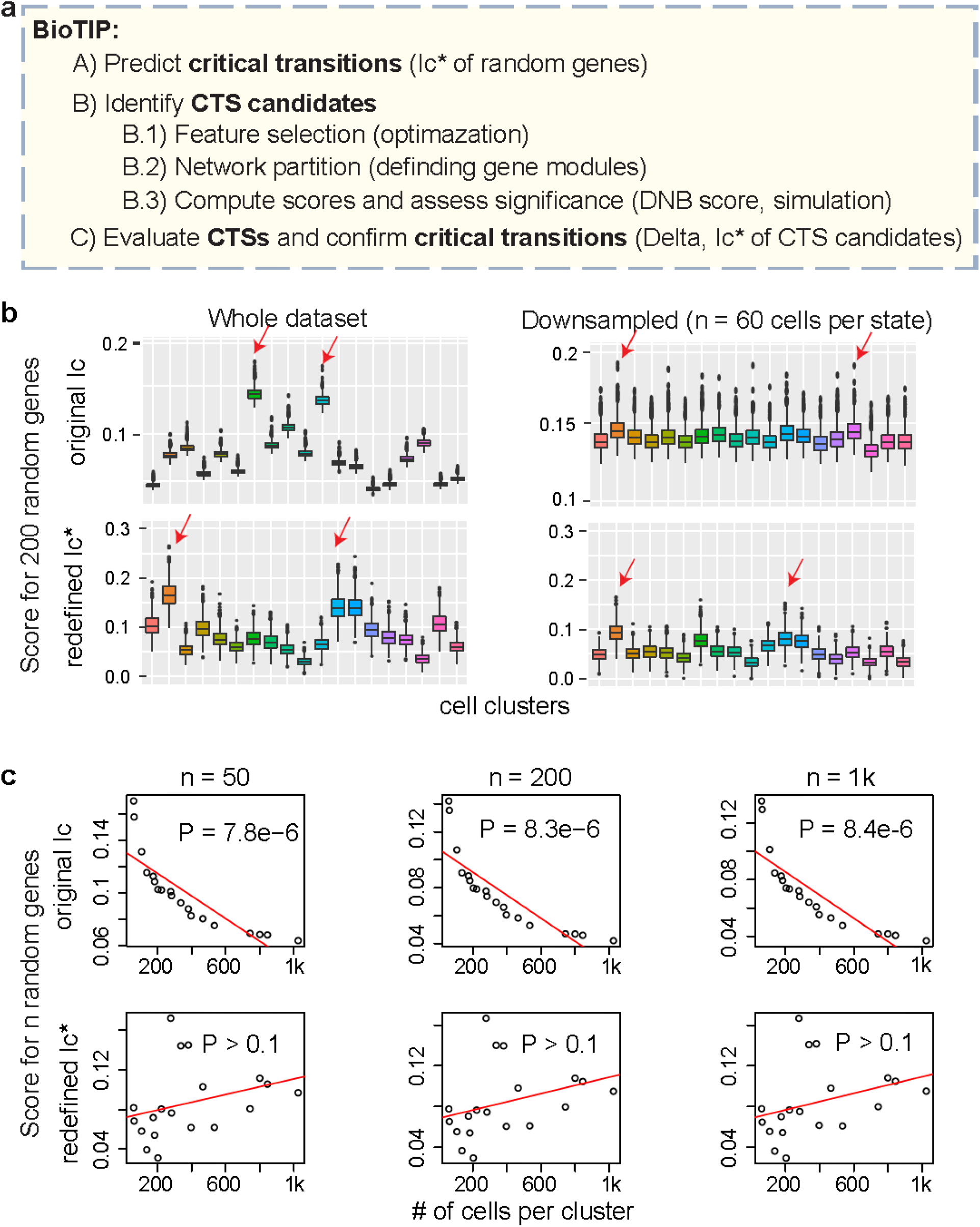
BioTIP workflow, in which Ic* yields consistent results to compare states with variable cell population sizes. **a**, BioTIP’s five-step analytic workflow. **b**, Comparison of the Ic scores on a simulated scRNA-seq dataset of variable subpopulation sizes (left, gene permutation based on the data of GSE87038) with that of down-sampled equal subpopulation sizes (right). Boxplot showing the simulation of 1000 scores based on 200 random genes. Box color decodes 19 unique cell clusters. In each subpanel, two red arrows point the top two scores. Ic scores output different clusters with highest scores (top); Ic* output consistent results (bottom). **c**, Calculating Ic-scores on the simulated dataset, using different numbers of random genes (50, 200, and 1000, respectively). Each dot represents one cell subpopulation (cluster). A p-value of Pearson Correlation is calculated between population sizes (x) and Ic scores (y) across all states, for which a linear regression line presents the strength and direction. Ic scores significantly decreases with large state sizes (top); Ic* does not significantly associated with state size (bottom).

1. higher DNB score at the state from which this CTS was identified, thus selecting CTS candidates
2. the highest Ic* at the same state, thus characterizing its indicative critical transition
3. significantly higher Delta at the same state than that of random genes, thus distinguishing the CTS from gene expression noise.

### 2.2. Ic* Detects Tipping Points More Accurately Than the Original Ic Scoring System

We first discussed two shortcomings of the existing Ic method for tipping-point identification using statistical-resampling based approaches. First, Ic calculated for each state as the ratio of average between-gene correlations to average between-cell coefficients and compared among states (Mojtahedi et al., 2016). The calculation of between-gene correlations over samples in each state biases Ic towards small states, and sometimes led to wrong predictions. It is because of the serious defect with the classical estimation of a correlation matrix from a state with few samples. This defect impacts Ic’s performance to compare large sates with small states. To address this defect, we redefined a new **Ic**\* score by a ‘shrinkage estimation of correlation matrix’ method (Schafer and Strimmer, 2005) (**Methods**).

We characterize Ic*’s advancement on tipping-point prediction in simulated single-cell transcriptomes, where the largest state size was seven times larger than the smallest state size (**S method, Section 3**) (Pijuan-Sala et al., 2019). We first applied the Ic method to the whole dataset, then repeated the same Ic analysis after down-sampling the data to equal-sized states. Unintendedly, the earlier full-sized analysis and the later equal-sized analysis yielded different results (**Fig 2b, top**). By contrast, and rigorously, Ic* predictions were consistent regardless of the inputted cell numbers per state (**Figs 2b, bottom**).

The advance of Ic* is universal, regardless of the number of genes tested (from 50 to 1000, **Fig 2c**). When calculated with 200 random genes, a positive correlation resulted between the Ic scores and the cell-population sizes (P= 8.3e-6). Although we expect critical transition states to be smaller than stable states in a snapshot data, this result is incorrect for the system because it was irreproducible in a down-sampled simulation (**Fig 2b**). By contrast, using Ic* to analyze the same dataset removed this correlation (**Fig 2c, bottom**). This suggests that when using Ic* instead of Ic, there is no longer a risk to find incorrect bifurcations when working with data sets or variable population sizes.

### 2.3. Applying BioTIP to benchmark dataset with an experimentally verified bifurcation and driving TFs

Applying BioTIP to single-cell gene expression profiles, we showed that it not only identified multiple bifurcations but also characterized them by significant CTSs. We demonstrated the accuracy with an experimentally validated bifurcation in early cardiogenesis, when induced pluripotent stem cells differentiating into cardiomyocytes (Bargaje et al., 2017). This dataset contains the gene expression profiles of 96 developmental genes for 1,934 cells collected from six timepoints (Bargaje et al., 2017) (**Fig 3a**). Day 2-2.5 was the verified developmental pitchfork bifurcation when multipotent primitive streak (**PS**)-like progenitor cells branched out into either the mesoderm cardiomyocyte (**M**) lineage (marked by Sox17) or the competing endoderm (**En**) lineage (marked by Hand1). **Fig S1a** presents the lineage mark gene expression.

**Figure 3.**
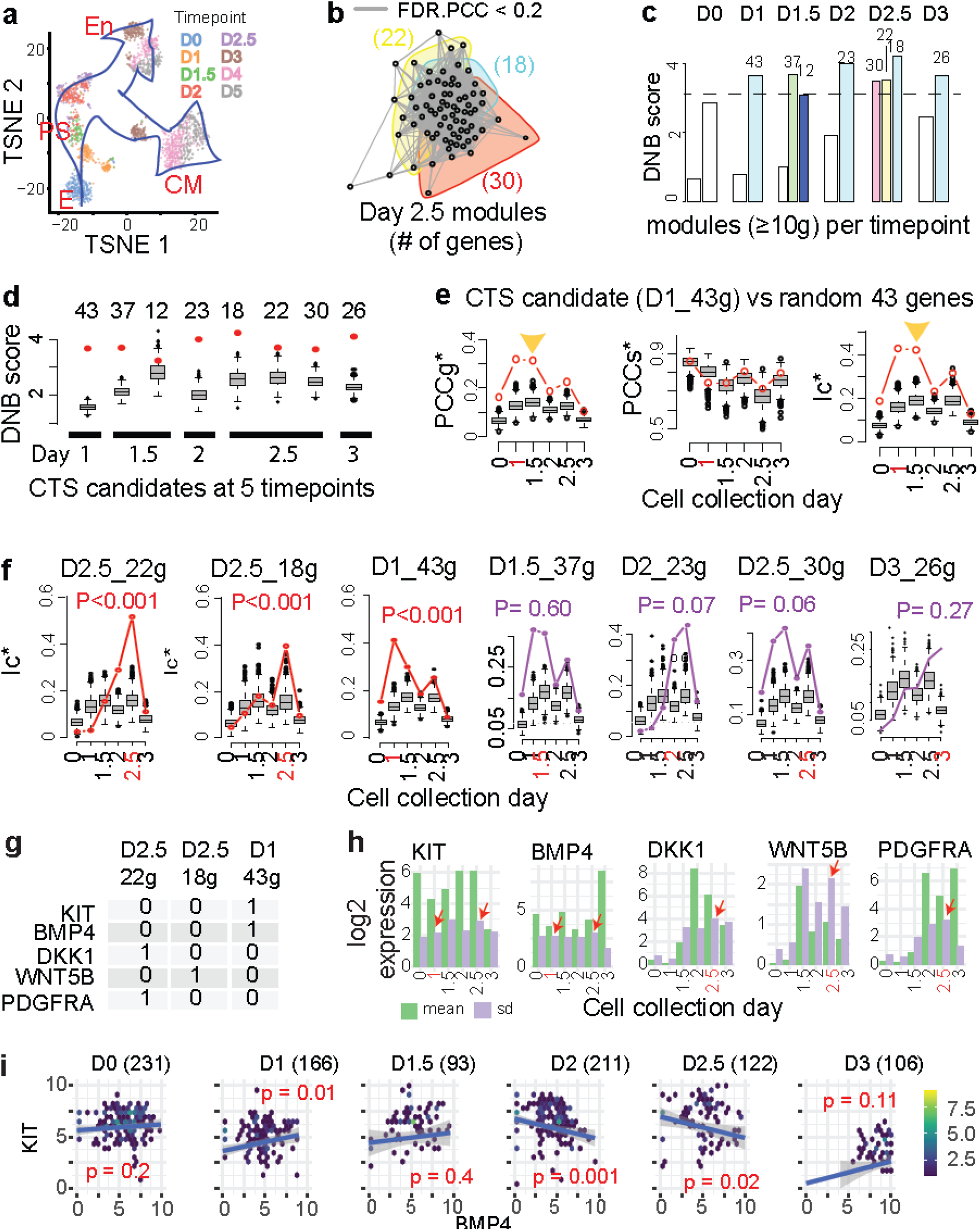
BioTIP identified multiple significant CTSs that contain experimentally validated key TFs controlling fate choices in early cardiogenesis. **a**, TSNE plot of the 1,896 induced stem cells, colored by 8 collection timepoints. Cell identities were supported by marker expression (**Fig S1a**). E: epiblast, PS: primitive streak; CM: cardiac mesoderm, En: endoderm. Hollow blue arrows illustrate the knowledge-based cell differentiation trajectories. **b**, Network view of three gene modules detected by the random-walk algorithm at day 2.5, respectively in colored circles. Each module contains the genes (nodes) that present pairwise between-gene correlation in expression (edges, FDR of Pearson correlation test < 0.2) among 122 day-2.5 cells. Gene numbers (≥10, given in the parentheses) are shown and coded by colors. **c**, Bar plots showing DNB scores in each cell collection day (D). 14 modules with the identification of 10 or more out of the 96 measured genes are shown. The horizontal dashed line indicates a cutoff, resulting in 8 prioritizing modules with the gene count per module listed atop. **d**, Boxplot comparing the observed DNB score (red dot) with the random scores for the 8 prioritizing modules, respectively, with the gene number atop. Each box presents the results of 1000 runs on randomly selected genes of the same size, over the cells at its derived timepoint. 7 out of 8 prioritizing modules were significant (P<0.001, when a red dot above its box), being the CTS candidates. **e**, Boxplot comparing the observed score with the random scores for the CTS candidate of 43 genes detected at day 1. Random scores were calculated based on permutation of gene labels. Showing are the between-gene correlation (PCCg*, left), between-sample correlation (PCCs*, middle), and Ic* score (right), respectively. Yellow triangles point the significantly high PCCS* and the Ic* at unintended day 1.5, respectively, suggesting these 43 genes are not day 1 specific. **f**, Ic* of 7 CTS candidates (red or purple line) compared to that of random-gene simulations (boxes, 1000 runs) across all timepoints. Left 3 panels are significant (P<0.05 at the predicted timepoint which is labeled in red on the x-axis); right 4 are insignificant, being the false positive. **g**, Table view of the presence (1) of five known cardiogenesis TFs in the three identified CTSs. **h**, Bar plot displaying the log2-scaled expression patterns of the five cardiogenesis TFs across timepoints. For each TF, red arrows point the highest standard deviation (sd) which is independent of the highest mean value. **i**, Hexbin plots comparing the average expression values of two TFs among cells collected at each day, respectively. Cell population sizes are given atop in the parentheses. A p-value of Pearson Correlation is calculated between BMP4 (x) and KIT (y), for which a linear regression line presents the strength and direction. In panels e. f and h, each CTS candidate-predicted timepoint is labeled in red on the x-axis.

We applied BioTIP to the temporal gene expression profiles of 929 cells (days 0 - 2.5, and mesoderm-specific day 3) as previously applied to Ic (Bargaje et al., 2017). We first partitioned all 96 genes into modules that highly co-expressed in cell at any given collection day (**Fig 3b**). Among the 8 prioritized modules (**Fig 3c**), we detected 7 CTS candidates at five timepoints (P<0.001, **Fig 3d**), based on the DNB scoring. Although a 43-gene module identified at day 1 present significantly higher than random Ic* scores at both days 1 and 1.5, due to their high between-gene correlations at both days (**Fig 3e**). Therefore, BioTIP reject his identification. Being evaluated by Ic* similarly, only three CTSs at days 2.5 or 1 were found to be significant (P of Delta < 0.05, **Fig 3f**).

The highest Ic* was observed at the verified bifurcation at day 2.5. By contrast, Ic had been applied to the same profile to assign an equally high score at days 1.5 and 2.5 (Bargaje et al., 2017), which is presented by two equally high simulation scores (boxes at days 1.5 and 2.5 in **Fig 3f**). The advantage of BioTIP over Ic is that BioTIP reflects the interaction patterns of module genes whose covariation increases before (or at) the critical transition subpopulation (Richard et al., 2016; Teschendorff and Feinberg, 2021).

We conclude that BioTIP analysis is more accurate and precise than existing methods -- having higher accuracy than Ic (which failed to prioritize day 2.5) and higher precision than DNB (which had false positive) in the prediction of tipping points. Additionally, applying BioTIP to single-cell gene expression can capture multiple tipping points. Furthermore, our data supports the theoretical hypothesis that CTSs can describe both topologically pitchfork (e.g., day 2.5) and saddle-node (e.g., day 1) bifurcations.

### 2.4. Significant CTSs Disclose Key Transcription Factors Controlling Cell Fate decisions

We collected five established cardiogenesis TFs in this system which are KIT, BMP4, DKK1, WNT5B, and PDGFRA (Bargaje et al., 2017). In this system, the stem cells were induced by BMP4 and Wnt pathway activator; KIT was the experimentally verified bifurcation predictive marker that guides differentiation into the mesodermal cardiomyocyte lineage as opposed to the competing endoderm lineage; And the expression levels of DKK1, WNT5B, and PDGFRA were highly correlated with the BMP-induced differentiation efficiency towards cardiac cell fate (Bargaje et al., 2017). All five key TFs were disclosed by the three computationally identified CTSs (**Fig 3g**).

The identification of these key TFs is unique, being overlooked by conventional group-mean based approaches. On one hand, these TFs gained expressional variance at the bifurcation state (i.e., increasing standard deviation (sd), **Fig 3h**, purple bars). On the other hand, it is a set of genes’ covariations rather than single gene’s expression that characterize a bifurcation. For example, KIT and BMP presented increased co-expression at days 1 and 2-2.5, the identified and verified tipping points in this system (**Fig 3i**).

We conclude that BioTIP is a powerful approach to disclose key TFs at differentiation bifurcations by the identification of significant CTSs.

### 2.5. Applying BioTIP to Cell Clusters Identify CTSs as Key TF-regulated Targets

We propose a better detection of key TFs when applying BioTIP to cell clusters defined by the similarity of gene expression profiles. To test this hypothesis, we reanalyzed the same 1,934 cells that were previously grouped into 19 consensus clusters (Bargaje et al., 2017) (**Fig 4a, left**). Note that cells collected at the same timepoint can come from distinct cell clusters (**Fig 4b**), timepoint-based CTS identification may describe the heterogeneity regarding multiple clusters collected at the same day. To compare with the above time-course analysis, we applied BioTIP to nine clusters of the analyzed 929 cells.

**Figure 4.**
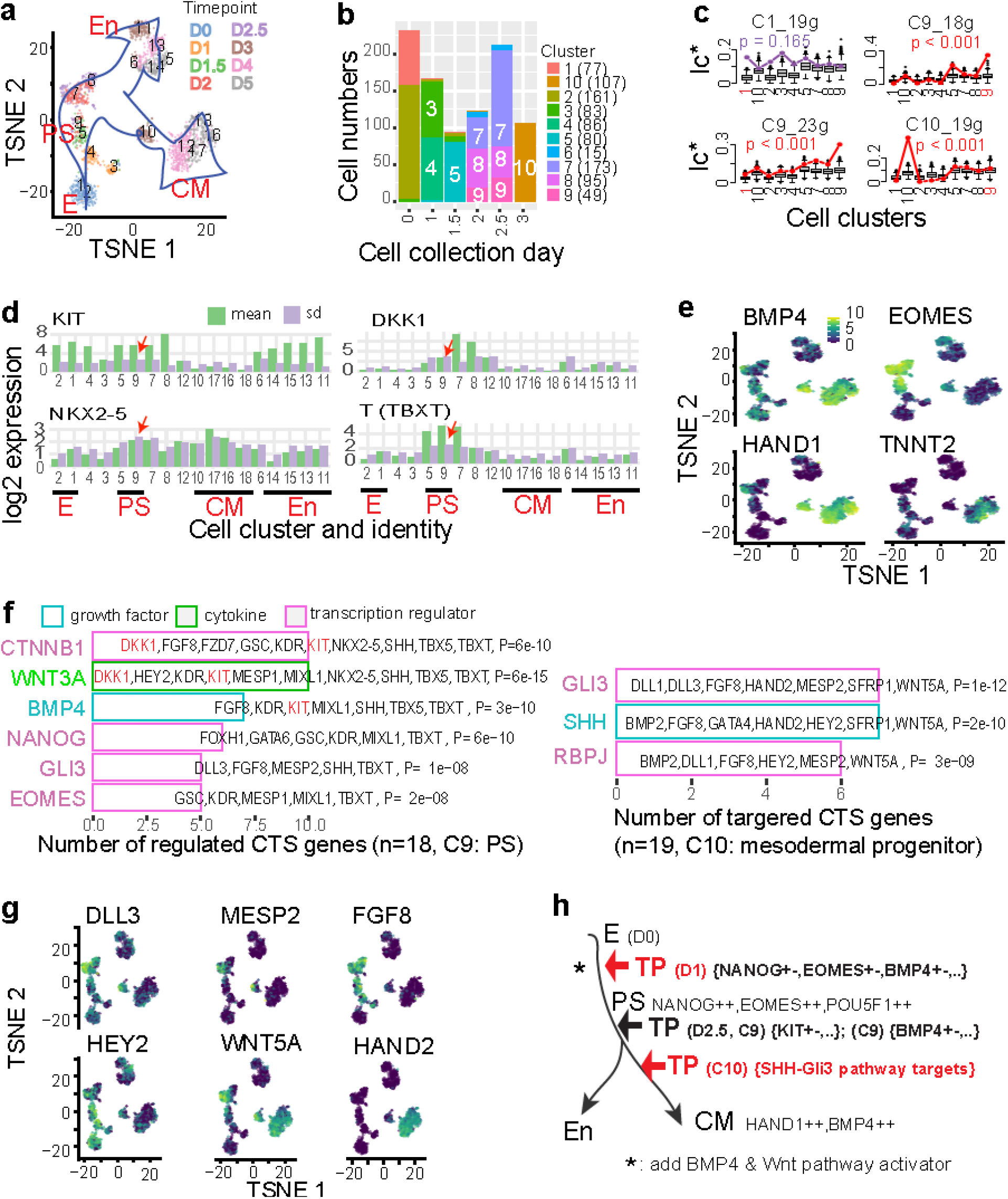
BioTIP identified significant CTSs that are independent of differentiation trajectory topologies and enriched for targets of fate decisive TFs. **a**, TSNE plot for all collected cells. Cells are labelled by 18 unique clusters of similar gene expression patterns (left). **b**, Stack bar chart projecting cell clusters into their collection days of interest (days 0, 1, 1.5, 2, and 3/M – only mesoderm-specific cells). Cell numbers are given in the parentheses (n=929 in total). Note that there are small proportion of endoderm-specific cells collected at day 2.5 (cluster 6, n=15). **c**, Similar to **Fig 3f** but showing the Ic* scores across cell clusters. For each of the four DNB-identified CTS candidates, also shown is the p-value of Delta scores at the predicted bifurcation state (cluster), resulting in 3 significant CTSs and one false positive. **d**, Bar plot displaying the log2-scaled expression patterns of four CTS member genes for the C9. For each gene, red arrows point the highest standard deviation (sd) exhibiting at C9. **e**, TSNE plot for all collected cells. Cells are colored by mark gene expression levels. **f**, Bar plot of significant upstream regulators for the 18 genes charactering C9 (left), or for the 23 genes charactering C10 (right), respectively. Also shown are the target genes and enrichment p-values (IPA analysis). Bar color decodes the molecular types of these upstream regulators. The experimentally verified bifurcation-driving TF KIT is highlighted in red. **g**, TSNE view of six C10 CTS member genes that are enriched for the SHH-GLI3 singling pathway. **h**, Tipping points (TPs) along the trajectory from early stem cell to CM differentiation, induced by BMP4 and Wnt pathway activator. Each cell state (stable or transitional) can be marked by specific transcription factors. Color distinguishes the verified tipping point and factors (black) or the BioTIP-identified new ones (red). The verified tipping point at day 2.5 was identified by both time-course and cellular cluster analysis using BioTIP.

Among four DNB-identified CTS candidates (**Fig S1, b-d**), we verified the significance of three CTSs, respectively indicating critical transition at clusters 9 and 10 (**Fig 4c, Table S1**). Cells of cluster 9 (C9) were collected at Day 2-2.5 (**Fig 4g**), the experimentally verified bifurcation of this dataset (Bargaje et al., 2017), and presented an abundant level of NANOG and EOMES representing the transient PS state (**Fig S1a**). Cells of cluster 10 (C10) were collected at day 3, when major lineage commitments took place (Bargaje et al., 2017). Cells in this cluster presented high expression of the mesoderm markers HAND1 but not the cardiac mark TNNT2 (**Figs 4e, S1a**), suggesting a mesoderm progenitor state. However, this cluster was overlooked in the original study (Teschendorff and Feinberg, 2021). Because C10 sits along the path towards mesoderm lineage, being a topologically a saddle-node bifurcation, it is also undetectable by other existing methods. Therefore, the identification C9 demonstrates the accuracy, but C10 the sensitivity, of BioTIP application.

We identified two distinct CTSs for cluster 9, with 18 and 23 genes in each. The CTS of 18 genes contained not only the verified bifurcation mark genes KIT and DKK1 but also the cardiac markers NKX2-5 and TBXT. These TFs gained variance in expression among C9 cells (**Fig 4f**, purple bars). These results show that BioTIP analysis can discover key TFs with increased fluctuation at bifurcation state, being potential cell engineering targets.

Our results further witnessed a model of CTS to be strongly enriched for direct and indirect targets of key TFs controlling fate decisions (reviewed in (Teschendorff and Feinberg, 2021)). Along the differentiation from PS to mesoderm, the 18 CTS genes of C9 were the enriched targets of PS markers NANOG and EOMES as well the cardiac mesoderm inducing marks BMP4 and WNT3A (IPA analysis, P< 1e-8, ≥5 genes, **Fig 4f, left**). Another set of 23 CTS genes of C9 were the enriched targets of not only BMP4 but also the pluripotency factor POU5F1 that is highly expressed in the primitive streak (Mohammed et al., 2017). These results fit the hypothesis of opposing factors causing transcriptional priming (Ranzoni et al., 2021; Teschendorff and Feinberg, 2021) (**Fig 1a**). Therefore, tipping-point theory based BioTIP analysis can discover targets of these key TFs.

Similarly for C10, the 19 CTS genes include the Wnt signal mark WNT5A. WNT5A level was induced at PS and maintains until cardiac mesoderm (CM) clusters (**Fig 4g**), agreeing with the literature that BMP4-Wnt activity within cardiac mesoderm appears to be positively amplified early commitment steps to cardiomyocyte have occurred (Kwon et al., 2007). Three upstream regulators of the 19 CTS genes were Sonic hedgehog (SHH), SHH downstream factor GLI3, and RBPJ (**Fig 4f, right**). Inhibition of these three factors has been shown to improve specification of cardiomyocytes (Diaz-Trelles et al., 2016; Parikh et al., 2015). We thereby reason that SHH-GLI3 cascade signal is one driver of the bifurcation at C10. Indeed, the expression levels of GLI3 targets DLL3, MESP2, and FGF8 are higher in earlier primitive streak (PS) state, as a readout of the decreasing attractors. In contrast, HAND2 is more highly expressed in later CM clusters, as a readout of an emerging new attractor. These patterns support the notion that saddle-node bifurcations can be seen as a gradual destabilization of previously stable attractor states, and the simultaneous emergence of new differentiated states (Teschendorff and Feinberg, 2021) (**Fig 1a**).

Overall, BioTIP analysis on cell clusters of distinct expression patterns found verified key TFs and disclosed new TFs (or signals) for further investigation. We show the success in identifying significant CTSs in two types of bifurcations, independent of observed trajectory topologies, in two ways (**Fig 4h**). First, the TFs driving cell differentiation from one stable state to a later state could gain its expression level with fluctuance, being not only a CTS member gene but also the marker of the coming state. Second, the competing regulators of the early and later states cause fluctuance in their target genes, allowing CTS to infer key TFs from enriched upstream regulators. We propose that BioTIP analysis of single-cell gene expression profiles can detect critical cellular state bifurcations when external signaling can redirect the population’s lineage fate.

### 2.6. Application to Mouse Gastrulation Identifies Four Robust CTSs

To further demonstrate applicability, we studied mouse embryonic developmental bifurcations. We hypothesize that BioTIP analysis could capture the regulated stochasticity and reproducibly indicate bifurcations in independent samples on the same developmental paths. To test this hypothesis, we studied a mouse gastrulation *in-vivo* dataset (Pijuan-Sala et al., 2019), focusing on early organogenesis at embryonic day (E) 8.25 when precursor cells of major organs have been formed (Kaufman and Bard, 1999). We clustered 7,240 E8.25 cells into 19 states (sub-populations) and annotated them by up-regulated transcriptional markers (**S Methods**). The branches of cardiac mesoderm and muscle mesenchymal from early mesoderm were followed by a bifurcation between the hematopoietic and endothelial lineages (**Fig 5a**). Note that S13, S10, and S15 were three haemato-endothelial progenitor states sequentially in pseudo-order. They also have notable differences in gene expression: the early S13 has the highest average expression of *Kdr* (**Fig S2a**); S13 and S15 were closely clustered groups (positive silhouette width) but multiple clusters intermingled at S10 in expression space (**Fig S2b**), suggesting branching events at S10.

**Figure 5.**
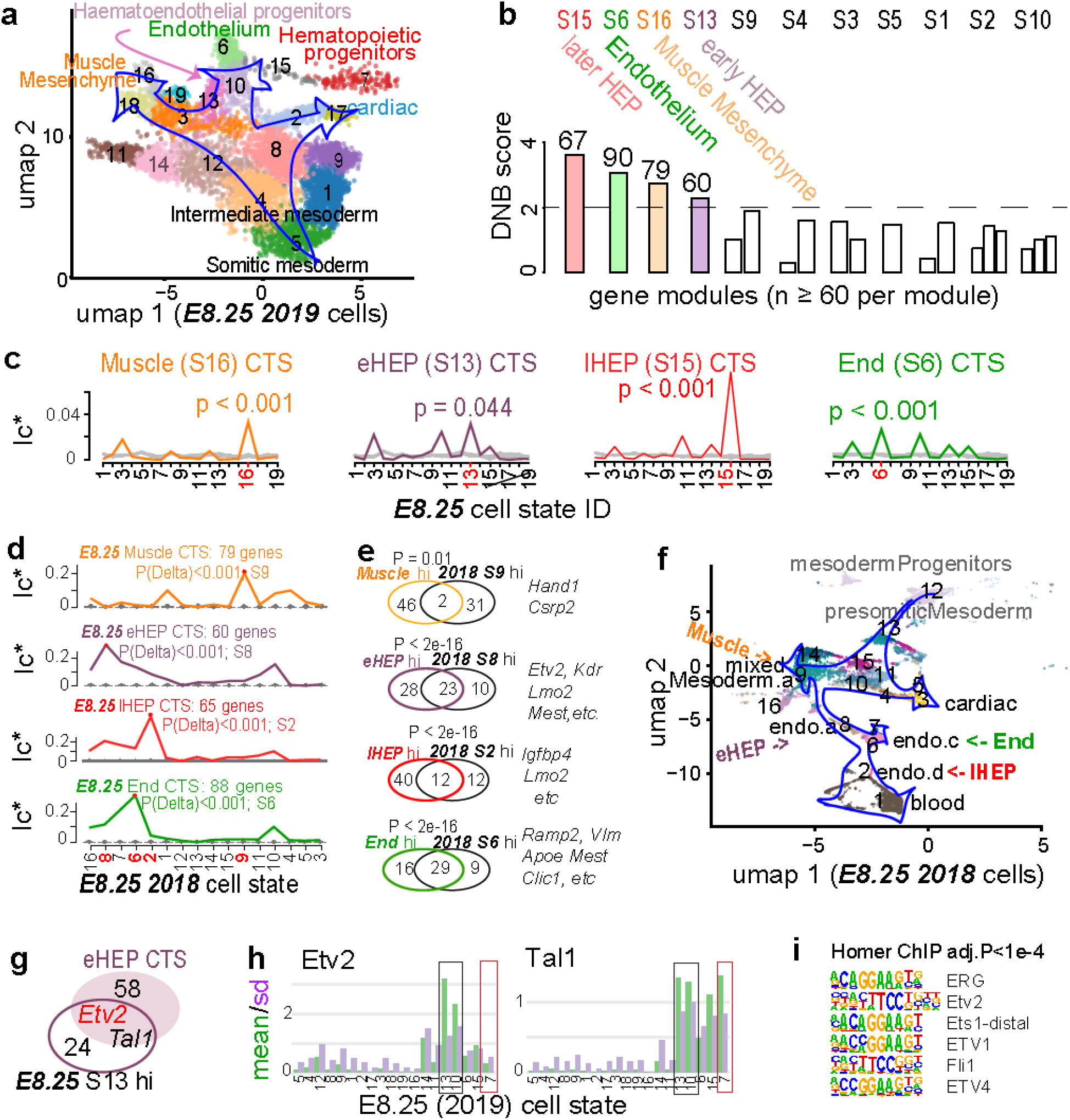
Applying BioTIP to scRNA-seq data of developing mesoderm disclosed the cell-fate decisive ETV2 and revealed reproducible CTSs. **a**, Uniform manifold approximation and projection (UMAP) plot of 7,240 developing mesoderm cells (E8.25, published in 2019), colored and numbered by 19 unique clusters (states) of transcriptionally similar cells. Blue hollow arrows illustrate the knowledge-based pseudo-orders. **b**, Bar plots illustrate the DNB scores in each E8.25 state (S). The horizontal dashed line indicates a significance threshold a cutoff, resulting in 4 prioritizing modules with the gene number atop. Eleven states with module identifications of 60 or more genes are shown. **c**, Evaluation of four identified CTS candidates, each being labeled by the cell-cluster identity from which the CTS was identified. For each CTS candidate, comparing its Ic* (colored line) to the gene-size-controlled random scores (light grey lines, 1000 runs). Also showing is the p-value of Delta of the observed Ic* at the CTS-indicated tipping point. **d**, Validating identified CTSs in independent profiles of 11,039 E8.25 cells (published in 2018). Ic*s of each CTS across new cell subtypes (line) were compared to their empirically simulated scores (box, 1000 runs). Calculation was conducted after mapping all CTS genes (n=67, 90, 79, 60, respectively) to this profiling, extracting above 98% of CTS genes (n=66, 88, 79, 60, respectively) measured. For each CTS, the subtype with the highest Ic* is highlighted with a red dot, and the p-value for the Delta at that point is shown. The red font on the x-axis indicates ‘CTS-mapped’ states of bifurcation. **e**, Venn-diagram comparing the up-regulated biomarkers of each bifurcation state (in the 2019 profiles) with the biomarkers of the mapped cell subtypes (in the 2018 profiles). Circle color decodes the four CTSs. The numbers in the Venn count the biomarkers. OR: Odds ratio. Fisher’s Exact test. Some shared marker genes are displayed to the right. **f**, UMAP showing the 2018 E8.25 cells *in vivo*, colored and indexed by 16 unqiue subtypes. Blue hollow arrows illustrate the knowledge-based pseudo-orders. Colored text indicates four tipping points mapping to four subtypes. **g**, Venn-diagram comparing the E8.25 early HEP (eHEP) CTS genes and up-regulated biomarkers of this cell cluster. **h**, Bar plot displaying the log2-scaled expression patterns of Etv2 and Tal1 across cell clusters. Colored squares highlight the two HEP (black) or HP (red) clusters, respectively. **i**, Significantly enriched ETS-binding motifs found by Homer from the promoters ([-200, 100] around transcript start sites) of the 60 CTS genes. **j**, Venn-diagram showing the overlap among E8.25 eHEP CTS genes, the CRISP-validated upstream regulator of Etv2, and the Etv2 direct targets in a previous ChIP-seq experiment. **k**, A model of Etv2 masters the HE bifurcation. The yellow ball is the initial state. As Etv2-targets, increasing fluctuation in CTS-gene expressions (pink) can be triggered by Etv2 autoregulation. After entering later stable states, some oscillated CTS genes gain enriched expression.

Applying BioTIP to all 19 cell states regardless of their pseudo-trajectory, we detected four CTS candidates. Each presented significantly high DNB scores at one cell state, respectively (**Figs 5b, S2c**). The significance of these CTS was confirmed by Ic* simulation (P<0.05, **Fig 5c**). These CTSs revealed four transcriptional priming states, respectively (**Tables 1, S2– S5, Fig S3a**). Along the trajectory from early mesoderm to endothelium, there are many tipping points when a cell transitions from one stable state to another. Therefore, BioTIP’s ability to identify multiple significant tipping points is essential to understand the dynamic processes in development.

**Table 1.**
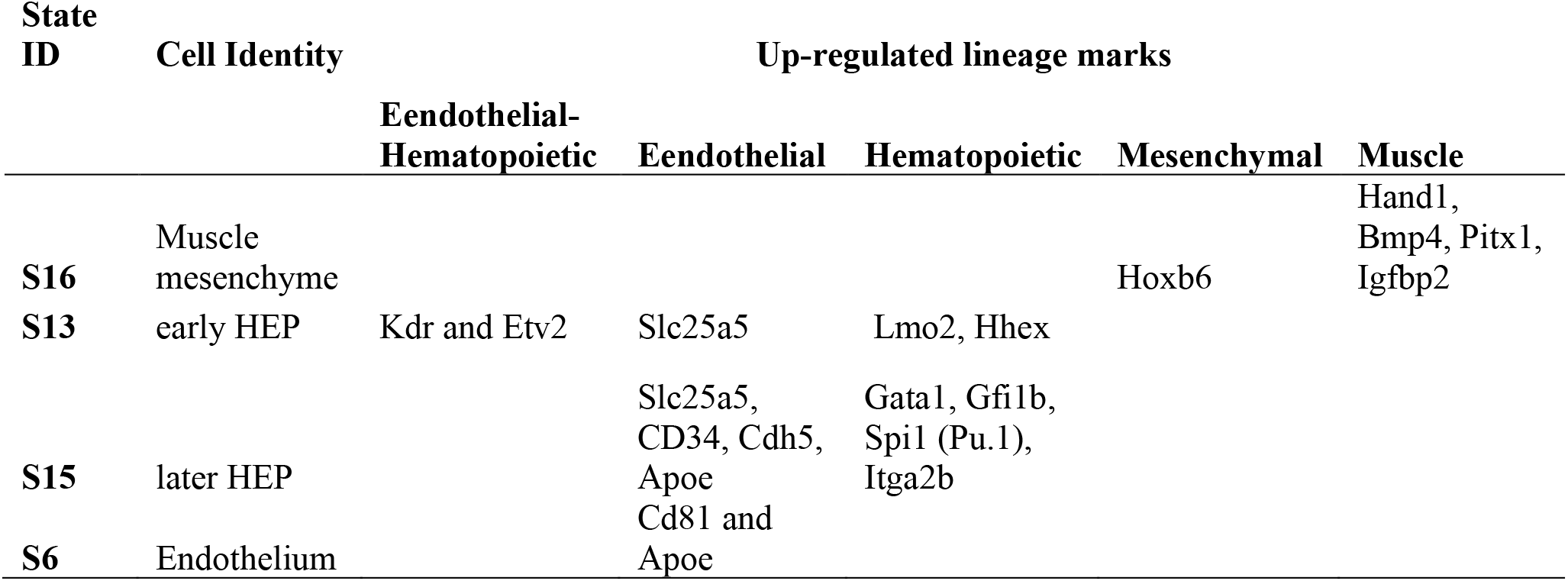
The established cell lineage marks.

Given a significant CTS, its significance in indicating a critical transition should be concordant in independent profiles of the same developmental stage. We tested this hypothesis using *in vivo* samples (E-MTAB-6153, published in 2018) (Ibarra-Soria et al., 2018). We reanalyzed 16 previously defined subtypes of E8.25 developing mesoderm (11,039 cells) that spanned from mesoderm progenitor to blood and endothelium.

Of each our identified CTSs, the Ic* scores over 19 cellular subtypes peaked at one subtype significantly (P of Delta<0.001, **Fig 5d**), thus pairing each detected bifurcation sate to predefined subtypes. The agreement between each pair was supported by their shared biomarkers (**Fig S3**). Three out of four pairs shared the top-10 up-regulated biomarkers significantly (12-29 shared markers from 10k common background genes, Fisher’s exact test P<2e-16, odds ratio> 250, **Fig 5e**). One pair with moderate overlap (P=0.01, OR=15) was between the muscle mesenchyme specification and the ‘mixed mesoderm subtype a’, sharing the established cardiac mesoderm marker Hand1 and cell growth regulator Csrp2.

We then verified the agreement between each pair in development pseudo-order. All four CTS-mapped cell subtypes resembled the pseudo-order of the four CTS-indicated bifurcations (**Fig 5f vs 5a**) – The muscle mesenchyme specification from the mesoderm was followed by the commitment to the early HEP state; then a transition into the later (more blood-committed) HEP state occurred before the specification to either blood or endothelium. These concordances support the significance of the four CTSs, suggesting they are robust in charactering bifurcations in developing mesoderm.

In summary, we identified four significant CTSs as the readout of imminent developmental bifurcations and evaluated their reproducibility in independent samples. This novel angle of transition states reflects the plastic and combinational regulatory mechanisms regulating early mesoderm development.

### 2.7. CTS-based Mapping of Developmental Bifurcations

We have shown that significant CTS contains or is the targets of key TFs marking or driving the bifurcation. Here, asked whether the identified CTSs disclose fate-decisive key TFs. From four identified CTSs we isolated three TFs whose fluctuation in expression most likely impacts other CTS genes’ fluctuation; these TFs were Gata1, and two ETS family members Ets1 and Etv2 (**Fig S2d**). Each TF was not only the CTS member gene gaining covariance with other CTS genes but also the upstream regulator whose variance could explain the expression changes of the CTS genes (**Methods**). Among these three TFs, the CTS member gene ETV2 also presented as traditional up-regulated marker for its indicated cell state S13 (**Fig 5g**), suggesting its transcriptionally regulatory role.

To verify the CTS-driving role of Etv2, we checked two key ingredients of a bifurcation (Scheffer et al., 2012): threshold-dependence and autoactivation. First, the variance of *Etv2* expression peaked at HEP before HE bifurcation, suggesting that *Etv2* fluctuation marks this fate decision; while *Etv2* reached its highest mean value at early HEP, indicating an expression threshold is required (Koyano-Nakagawa and Garry, 2017; Zhao and Choi, 2017) (**Fig 5h**). By contrast, the variance and mean value of *Tal1* expression peaked at early HEP and then maintained its value for blood progenitors, going along with literature that a positive Etv2-Tal1 loop is induced upon the activation by *Etv2* (Zhao and Choi, 2017). Second, we observed enriched ETS motifs, denoting Etv2 as an upstream regulator of this CTS (**Fig 5i**). Consistent with this observation is the significant overlaps between 60 early HEP CTS genes and the ChIP-seq detected Etv2 targets in hematopoietic and endothelial lineages (Liu et al., 2015) (P=3.8e-7, **Fig 5j**). Meanwhile, 60 CTS genes significantly overlapped with the upstream regulators of Etv2 during haemangiogenic differentiation (Zhao and Choi, 2017) (P=3.2e-11, **Fig 5j**), suggesting Etv2 is also a downstream target of this CTS. These results agree with reported Etv2 autoactivation in hematoendothelial specification (Koyano-Nakagawa et al., 2015). Therefore, we infer a bifurcation-driving role of Etv2. This inference agrees with literature that Etv2 activation specifies HE lineages and is required until HE bifurcation (Koyano-Nakagawa and Garry, 2017; Liu et al., 2015).

We further inferred the Etv2-mediated gene regulations from the Etv2-centered E8.25 60 CTS genes, which revealed four co-regulators (as both regulators and targets) -- *Tal1, Lyl1, Rhoj* and *Rasip1* (**Fig 5j**). Existing literature supports a strong cooperation between the Etv2 targets *Tal1* and *Lyl1* during HE transition (Guibentif et al., 2017), and both CTS members *Rasip1* and *Rhoj* are endothelium-specific Etv2 targets (Palikuqi et al., 2020; Singh et al., 2020). Given the consensus that VEGF signaling plays an instructive role in promoting *Etv2* threshold expression (Zhao and Choi, 2017), we model an Etv2-induced HE bifurcation (**Fig 5k**). In this model, positive feedback loops amplify the perturbations of a Etv2-activated GRN module, followed by negative feedback loops that repress *Etv2* and drive the cells into distinct differentiated, stable states. Together, this is the first computational inference for the transient Etv2 expression that is required for HE bifurcation.

Similarly, the predicted roles of Gata1 and Ets1 in marking or driving the bifurcation at later HEP are new, and our computational inference is partly supported by literature: Gata1 has been reported to drive stem cells toward erythroid/megakaryocytic differentiation (Grass et al., 2003). Ets1 has been shown to function redundantly with Etv2 in promoting embryonic vasculogenesis and angiogenesis (Casie Chetty and Sumanas, 2020).

In summary, our efforts to detect significant CTS from a noisy background moves the field one step closer towards understanding the dynamic TF regulation, in which the transient induction of a master TF concurs with critical state changes. The above results provide an impetus to adopt tipping-point analysis to uncover the roles of transcriptional mosaicisms with high resolution.

## 3. Discussion

In this paper, we detail BioTIP, a new workflow that advances mathematical tipping-point methods to identify significant CTSs. When the system approaches a critical state transition, the underlying gene expression signals become more stochastic, priming new states with distinct expression patterns. BioTIP parses out noisy transcriptional background signals and identify multiple significant, robust, trajectory topology-independent CTSs. Detecting these CTSs in cellular development is of prime interest for forecasting cellular engineering potential (Bargaje et al., 2017). Built on previous mathematical studies, BioTIP achieves three methodological contributions to study TF-regulated CTSs using scRNA-seq data:

1. BioTIP enables the identification of multiple CTSs from stochastic background signals. The false positives were removed in multiple simulation steps (rather than simply taking the maximum), particularly by inputting DNB-identified gene signals into to the Ic*-scoring system. Further advancements in the inputs of the DNB span the identifications. Consequently, the identified CTSs inform key TF regulations that were previously unable to computationally explore.
2. BioTIP explicitly compares Ic* scores over states respite population sizes to infer tipping points accurately. By an advanced estimation of correlation matrices (Schafer and Strimmer, 2005), BioTIP reduces the sample-size related bias in the CTS-evaluation step.
3. The focus on critical transitions is novel in current scRNA-seq data analysis. BioTIP enables the identification of CTSs from noisy data, which can be independent with trajectory topologies, enabling the CTS identification for potential saddle-node bifurcations.

Using BioTIP, we were able to computationally capture hematopoietic-endothelial bifurcations in gene expression changes, through which Etv2 expression is induced, fluctuating, and extinguished rapidly (Koyano-Nakagawa and Garry, 2017; Koyano-Nakagawa et al., 2015). Four CTSs were either distinguished from or recaptured 8-21% of the up-regulated marker genes, respectively, at a bifurcation state (**Fig S2e**), confirming the independence between CTS and differentially-expressed genes (Liu et al., 2019). This independence suggests that BioTIP analysis detects new features of dynamics, going beyond traditional analysis relying on differential expression patterns. BioTIP models the lineage specifications to be not just a simple gradual silencing of the progenitor cells with simple upregulation of lineage marker genes (Bargaje et al., 2017; Zhou et al., 2019). Such analyses allowed for Etv2 to be identified as a CTS driver and for regulatory loops it’s involved to be characterized. These findings demonstrate how BioTIP’s ability to identify bifurcations has applications across different types of transcriptome data and fields of biology.

BioTIP is based on the regulated stochasticity model. Indeed, cells are most likely to determine their fate by taking developmental cues from stochastic fluctuations. To the authors’ knowledge, BioTIP is the first workflow to correctly identify tipping points and significant CTSs despite the presence of stochastic fluctuations in expression (background gene signals). In the cardiogenesis data that random gene signals can be indicative of a tipping point, Ic* was able to capture significant CTS (**Figs 3-4**), suggesting that stochastic perturbations coexist with transcriptional priming in these systems. In the developing mesoderm data (**Fig 5**), the results show that a perturbation in CTSs (rather than expression noise) is the only readout of critical transitions. Knowledge-based TF-binding metrics help discriminate between TF-driven perturbations triggering bifurcations versus those perturbations responding to non-transcriptional stimulus. To unlock those mechanisms besides of or coexisting with TF-driven perturbations, future work will be a comprehensive integration of multi-omics profiles (Stuart et al., 2019; Su et al., 2020).

Existing biological tipping-point studies have not produced software packages. The only available tool, *earlywarnings* (Dakos et al., 2012) (https://early-warning-signals.org), has been evaluated and accurately predicts the epithelial–hybrid-mesenchymal determination (Sarkar et al., 2019). However, it only allows for univariate data analysis. Few scRNA-seq analytical methods can predict general phenotypic transitions without checking variance and correlation of gene expression. These methods include pseudo-ordering analysis of cellular states (Lummertz da Rocha et al., 2018; Wang et al., 2019), single-cell clustering using bifurcation analysis (Marco et al., 2014), unstable-cluster marking using the silhouette-width method (Rousseeuw, 1987), and cellular entropy changes (Mojtahedi et al., 2016; Teschendorff and Enver, 2017). Importantly, these methods cannot identify CTSs. Beyond all existing methods, BioTIP represents a new multivariate tipping-point analytic tool for the computational biology community.

Note that the BioTIP approach cannot distinguish pitchfork or saddle-node bifurcations and is not universally applicable. First, this work only detects tipping points based on certain early-warning signs such as increased variance and correlation, but there are many other warning signals associated with other types of tipping points. Second, transcriptional CTS cannot be significant when the noise in gene expression is larger than biological CTS (Oku and Aihara, 2018). Additionally, attractors are features of dissipative dynamical systems, but not all biological systems are dissipative.

In conclusion, we have developed the BioTIP algorithm and tool to identify CTSs from noisy gene expression profiles. We illustrate the utility and efficiency of BioTIP across multiple profiling technologies, system levels, and biological contexts. We suggest the use of transcriptomic tipping-point analysis for a deeper understanding of the plasticity, heterogeneity, and phenotypic changes in dynamical biological systems.

## 4. STAR Methods

### 4.1. Notation

Let *X* denote the *p* × *n* matrix of the expression levels of *p* genes ({*g*_1_, *g*_2_, … *g*_*p*_}) in rows and *n* samples ({*s*_1_, *s*_2_, … *s*_*n*_}) in columns. When the *n* = *n*_1_ + ∆ + *n*_*R*_ samples are divided into *R* distinct states, we can group the columns of *X* to have *X* = [*X*^1^|∆ |*X*^*R*^], where *X*^*r*^ denotes the *p* × *n*_*r*_ submatrix for samples in a state *r* ∊ {1,2, … , *R*} and *n*_*r*_ denotes the number of samples observed in the r-th state.

Given a state *r*, let 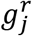 be the expression vector for gene *j* among samples in the state (i.e., the *j*-th row of *X*^*r*^), and let 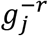 be the expression vector for gene *j* among samples outside the state (i.e., the *j*-th row of *X* where we remove *X*^*r*^). For a group of *q* genes (called a ‘**module**’) indexed by *m* = {*m*_1_, *m*_2_, … *m*_*q*_} ⊂ {1,2, … , *p*}, let 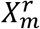 denote the *q* × *n*_*r*_ submatrix of *X*^*r*^ which selects the rows corresponding to the *q* genes in the module.

Given a gene expression matrix of interest *Z*, *PCC*_*g*_(*Z*) denotes the Pearson Correlation Coefficient matrix calculated between the genes (rows of *Z*), and *PCC*_*s*_(*Z*) is calculated between the samples (columns of *Z*) (i.e., *PCC*_*s*_(*Z*) = *PCC*_*g*_(*Z*^*T*^)).

Additionally, let ⟨·⟩ take the average of the off-diagonal entries in a square matrix, and let *sd*(·) be the operator that calculates the standard deviation of a vector.

### 4.2. Five steps in three components of the BioTIP workflow

**Component A: Predicting tipping point. Theory:** The original Ic calculated for each state as the ratio of average *PCC*_*g*_ (correlation coefficient matrix between genes) to average *PCC*_*s*_ (correlation coefficient matrix between samples) (Mojtahedi et al., 2016). Throughout states during a dynamic course, Ic peaks when the system approaches a critical transition because of the following two ‘early-warning’ features, exampled in the following single-cell scenarios:

a. The average *PCC*_*g*_ increases (**Fig 1a, S method, Section 1**). This is because *PCC*_*g*_ calculates the ratio of the between-gene variability to the average gene variability. Before transition, both variabilities are dominated by the noise around the attractor of the stable state. In contrast, when approaching transition, the decaying range-restriction effect limits the average gene variability within the unstable state – a decrease in the denominator of *PCC*_*g*_. Meanwhile, there is an onset of distinct states among the assembled cells. This new layer of fluctuations contributes to a higher between-gene variability – an increase in the numerator of *PCC*_*g*_. Together, *PCC*_*g*_ increases impending the transition.
b. The average *PCC*_*s*_ throughout the genes decreases. This is because the state of cell replicates become destabilized when approaching a tipping point, resulting in greater heterogeneity and thus weaker between-sample co-expression within the transition state.

**Problem:** The statistic 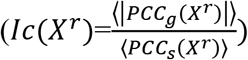 estimates the true correlation present in the data *X*^*r*^with the empirical correlation matrixes. Given a fixed number of genes *q*, the data varies in sample sizes {*n*_*r*_} among states; the numerator of Ic (each is calculated over one *n*_*r*_) exhibits serious defects with *X*_*r*_ that describes few samples (*n*_*r*_ ≪ *q*). It is because the empirical correlation matrix suffers from high variance, which inflates the numerator of the Ic for states with small sample sizes. Consequently, Ic has an undesirable performance on the comparison between *Ic*(*X*_*r*_): inaccuracy in its tipping-point prediction towards states *r* with a small *n*_*r*_.

**Method:** To address this problem, we first introduce a ‘shrinkage of correlation’ method to *PCC*_*g*_ (see (Schafer and Strimmer, 2005), the model named ‘Target D’ for details). This method shrinks the most extreme coefficients in the observed *PCC*_*g*_ towards more central value(s), thereby systematically reducing estimation error where it matters most (Ledoit and Wolf, 2004). Given a shrinkage target matrix *T*_*g*_,this method then weighs both matrices *T*_*g*_ and *PCC*_*g*_ to get an improved approximation 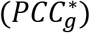 of the true (unknown) correlation matrix (**S Method, section 1**):

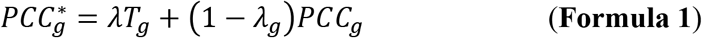

We set *T*_*g*_ to be the identity matrix -- so that we shrink the partial correlation coefficients in *PCC*_*g*_ towards 0 due to the globally low gene covariance in stable states; and the ones on the main diagonal represent the self-correlation of every gene.

The parameter *λ*_*g*_ ∈ [0,1] is a value to be estimated from *X*^*r*^ and *T*_*g*_, controlling how strongly the estimate 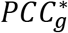 is shrunk towards *T*_*g*_; *λ*_*g*_ also controls the tradeoff between bias and variance when estimating the true correlation matrix (Schafer and Strimmer, 2005). When the sample size *n*_*r*_ is small, the optimal value for *λ*_*g*_ will be larger, leading to a stronger level of shrinkage of the partial correlations towards 0, which in turn shrinks the average of the *PCC*_*g*_ towards 0, limiting the effect of the problem mentioned above.

Similarly, we shrink the *PCC*_*s*_ to get an improved approximation 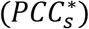 of the true (unknown) between-sample correlation matrix (see (Schafer and Strimmer, 2005), the model named ‘Target F’ for details). Particularly, we set the target matrix *T*_*s*_ with 1 on the diagonal and ⟨*PCC*_*s*_(*X*_*r*_)⟩ elsewhere, so that for each state, we shrink the correlations towards the average value. The choice of average not only reflects the steady expression pattern in a stable state but also preserves the pattern difference across states. Note that a different value of 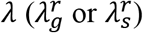 is estimated from the data when we calculate the 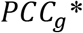 or 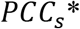 respectively for a state *r*.

*Step i)* **Finding tipping point.** We define a refined Index of critical transition (Ic*) scoring system, using this updated 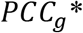 and 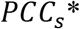 estimates:

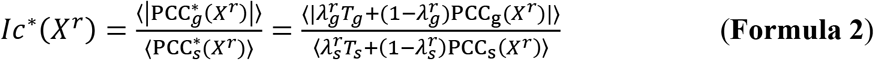

An increased 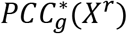, contributes to an increased Ic* regardless of sample size, and substantially enhances the reproducibility of the tipping-point identification from restricted sample numbers.

Simple algebra reveals a linear relationship between the original Ic and our improved 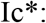 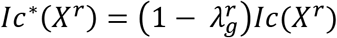 (**S Method, section 1**). As a sample size *n*_*r*_ raises, the calculatedvalue for 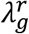 decreases to 0 and therefore we recover the traditional Ic score. Consequently, this correction enhances comparisons among states with variable sample sizes.

**Note:** Not all tipping points are predictable by Ic (using randomly selected genes) for which we document in this Result section. Therefore, the hybrid of Ic* with the following CTS identification is not trivial.

**Component B: Identifying significant CTS. Theory:** In the tipping-point model, each attractor (e.g., in the form of a fixed point along the dynamical trajectory) has a restricted effect (basin of attraction, e.g., expression regulation). The gradient of the basin of the attraction determines the rate at which a system turns back to the attractor from small perturbations. Thus, one indicator of tipping-point is a sudden ‘slowing down’ in which the system’s response to perturbations away from the current attractor decays just prior to the state transition (Scheffer et al., 2009). Not only is a tipping point characterized by an increase in *PCC*_*g*_ of a group of genes reflecting the onset of a new state, but their cell-to-cell variability also surge corresponding to the system’s loss of resilience (Richard et al., 2016). DNB has successfully quantified these characters from bulk-cell (Chen et al., 2012; Liu et al., 2019) also single-cell (Richard et al., 2016) expression profiles.

**Problem:** In previous DNB studies, significance, robustness, and reproducibility of the CTS identification are overlooked.

**Method:** Here, the goal is to identify a subset of genes that “drive” or respond to the state’s change at the tipping point. For expression profiles, we introduce new gene-feature selection, gene-module definition, and a module-size-adjusted DNB score (Chen et al., 2012) such that one can infer the underlying TF from the robust and significant CTS identification. With these refinements, BioTIP applies the concept of the DNB approach, by acknowledging its state-specificity.

*Step ii)* **Feature (gene) preselection.** The purpose of this step is to pre-select robustly informative transcripts and minimize the intrastate dispersions caused by sample outliers. This step is essential in high-throughput single-cell expression analysis as it picks the most informative genes from noisy background. To this end, we estimated a gene’s variation in state *r* relative to other states using a relative transcript fluctuation (**RTF**) score, for each state as follows:

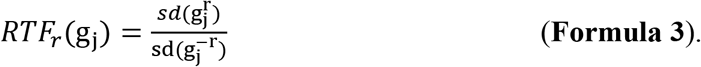

To ensure robustness to outliers, we used saturation (i.e., statistical-resampling) based approaches to optimize the RTF estimation. This optimization is done by 1) randomly drawing *n*_*r*_ × *b*% samples from a state *r* to estimate the *RTF*_*r*_(g_j_), and 2) repeating this drawing to select the *p* × *a*% genes that have top RFT scores. Parameter *b* is set to 80% to preserve the data structure, thus being optional and only recommended for large systems (*n*_*r*_>10 per state). By default, we set *a* up to 10% to focus on 5k or less genes with highest expressional fluctuation in a state (**Fig S1**, step 2.2). By adjusting *b*, we allow no less than about 200 genes per state for the following analysis. We document in the Result section that BioTIP is robust to this parameter.

*Step iii)* **Network partition**. Gene expressions are modulated by complex regulatory feedback loops; thus, the gene expression of a system can be conceptualized as an interconnected network. These networks are equilibrium for stable states -- if even one gene’s expression is perturbed, the effects could be kept in an attractor’s potential well throughout its connectivity in the network. When the perturbation passes a threshold of equilibrium, critical transitions happen (Scheffer et al., 2012). Therefore, it is essential to uncover which genes are most influential within these co-expressed networks or ‘**modules**.’ Decomposing pre-selected genes per state into several modules will serve as the inputs of the next CTS-searching step.

To identify significant CTS, it is necessary to partition genes into co-expressed modules. We chose to use random walk (RW) (Pons and Latapy, 2005) after reviewing an evaluation of 42 module-detection algorithms and finding RW to be the fastest, non-parametric, and have performed in the top 20% of the evaluated alternatives (Saelens et al., 2018; Wiwie et al., 2015) (**S Method, Section 2**). Additionally, BioTIP provides alternative methods such as k-means, HC, and PAM that showed superior performance in gene expression clustering (Wiwie et al., 2015).

*Step iv)* **Identifying non-random CTS**. We adjust the DNB’s bias towards small modules with a scaling factor equal to 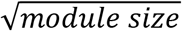, as being proposed recently (Yang et al., 2018). Comparing expressional deviation, the connectivity and homogeneity in this module relative to its complement set, and the scaling factor gives:

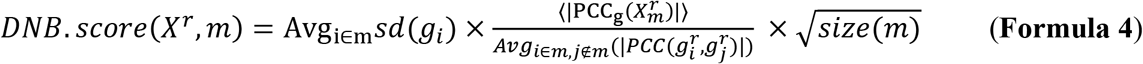

To identify the putative CTS, the modules 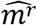 with the higher-than-expected DNB scores in the state *r*, we estimate random DNB scores from *X*^*r*^ by bootstrapping 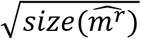 genes from the background.

**Three notes: 1)** An DNB-derived CTS is the genes members of 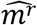 which are exclusively fluctuating and interactive in a tipping point *r*. **2)** When DNB works on the same networks preselected from the gene background, the state with highest DNB score indicates a tipping point state (Chen et al., 2012; Liu et al., 2017; Liu et al., 2019). This tipping-point prediction should agree with the prediction in Component A (if there is a prediction) because both methods identify increasing co-expression. **3)** The CTS-indicated tipping-point should be evaluable in the same data using different method, introduced in Component C.

**Component C: Two-way evaluation. Theory:** The early-warning features of a true tipping point should be captured by both Ic* and DNB scoring systems. With the same transcriptome data, we can evaluate the significance of the CTS and its indicated tipping point by the Ic* scoring system.

**Method:** The purpose is to define a tipping point by a significantly increased Ic* throughout discrete states. We design a Delta score to quantify abrupt changes in Ic* between states. Given a CTS candidate, the 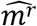 detected in state 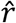, calculating its 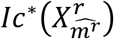 per state gives a vector 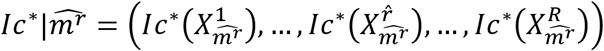, which should peak at the state 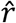 significantly. To quantify the module’s between-state changes, we propose a Delta score, the distance between the largest and the second-largest scores in 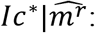:

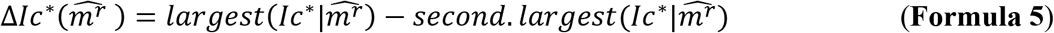

*Step v)* **Simulation study using Delta score**. Given a CTS identification, comparing the observed *Delta* score of the observed Ic* to the simulated *Delta* scores (of random Ic*s using randomly selected genes) gives empirical p-value. From the transcriptome *X*, the random Delta scores are calculated by fixing the state labels but randomly drawing the expressional values of *p* genes.

For a CTS-indicated tipping point state, comparing the observed *Delta* score to state-irrelevantly simulated *Delta* scores gives another empirical p-value. These random Delta scores are calculated by fixing the CTS genes but randomly assembling all *n* samples.

**Note:** In complex systems, the identified signal may be a mix of CTS along the trajectory of interest with unknown effects. Therefore, we recommend a validation using independent samples of concordant states on the same trajectory.

Finally, we comprise BioTIP into a comprehensive bioinformatics toolset for identifying transcriptomic tipping points and for describing critical transitions with non-random CTSs.

### 4.3. Prediction of upstream regulators

We predicted the upstream regulator for each set of the identified CTS genes using two strategies. First, we explained the expression changes of CTS genes with regulators whose change in expression are relevant to what is expected from the literature, using IPA (QIAGEN Inc., https://www.qiagenbioinformatics.com/products/ingenuity-pathway-analysis) (Kramer et al., 2014). The cutoff settings were FDR<0.005, at least 10% of target genes, and molecular type=’transcription regulator’. Second, because transcriptional regulation requires the binding of TFs, we searched for enriched ‘known’ TF-binding motifs which are mostly based on the analysis of public ChIP-Seq data sets, using Homer (Heinz et al., 2010). Significance settings were Benjamini-adjusted p<0.005 and at least 20% of target promoters ([−200,100] around TSS) with a known motif.

Motif enrichment analysis was also performed using Homer software (Heinz et al., 2010). Promoters were defined as a 200-nt window ([−200, +100]) around each TSS. These TSSs were directly extracted from the assembled transcripts of RNA-seq but were retrieved from the Ensembl (GRCm38_release97) annotations for the mouse gene symbols. Promoters overlapping with the blacklist were removed (Amemiya et al., 2019). We considered significance at a level of Benjamini-adjusted p<0.005 and at least 20% of target sequences with a known motif.

## 5. Data and Software Availability

The R package is available at https://github.com/xyang2uchicago/BioTIP.

Processed data and R codes are available at https://github.com/xyang2uchicago/BioTIP_application.

## 6. Acknowledgements

This work was supported by NIH grant 5R21LM012619 (XY, ZW), the University of Chicago: Biological Sciences Collegiate Division Quantitative Biology Fellowship (DG), Micro-Metcalf Program Project (YS), and Graham School of Continuing Liberal and Professional Studies, Master of Science in Biomedical Informatics Capstone Project (Biniam Feleke for building the Vignette, Qier An and Ieva Tolkaciovaite for testing functions for the initial package). We thank Bradley Leshem for scientific editing and Antonio Feliciano for testing the R package. We thank UChicago Center for Research Informatics for supporting high-performance computing services.

## 7. Author Contributions

XHY designed the study, managed the project, performed data analysis, contributed to the package, interpreted the results, and wrote the manuscript, with comments from JMC. ZW collected data, analyzed data, created the R package. AG contributed to the Ic* algorithm, method writing, and the initial R functions. YS wrote the online method part, updated the vignette, and maintains the R package. DA edited the manuscript and contributed to the initial scRNA-seq data analysis. MG and IM interpretated the results and revised the manuscript. YW wrote parts of the online method part.

## 10. Supplementary Figure Legends

**Figure S1.**
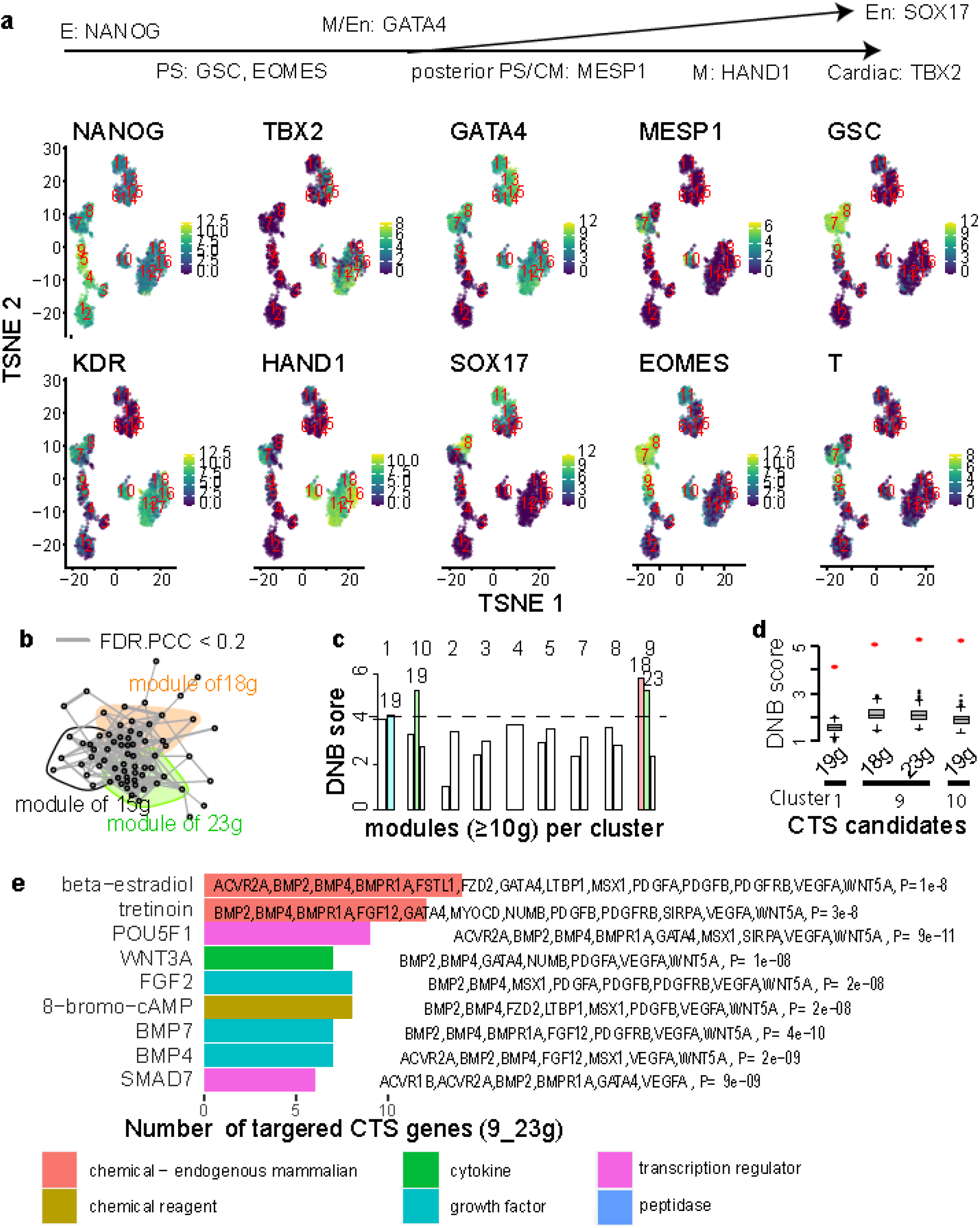
Analysis of single-cell RT-PCR data of induced stem cell towards cardio mesoderm. **a**, Top: Established cell lineage markers. Bottom: TSNE plot showing marker gene expression of individual cells, numbered by 18 unique cell cluster IDs. Each dot represents a single cell. Dot color decodes expression levels on log-2 scale. the E: epiblast, PS: primitive streak; CM: cardiac mesoderm, M: mesoderm, En: endoderm. **b**, Similar to **Fig 3b** but showing the three gene modules detected from cell cluster 9 (C9). **c-d**, Similar to **Fig3c-3d** but showing the DNB results of across cell clusters, resulted in 4 CTS candidates. Gene numbers are given at bottom. **e**, Bar plot of significant upstream regulators (left) for the 23 genes (right) charactering cluster 9 (C9). Color decodes the molecular types of these upstream regulators.

**Figure S2.**
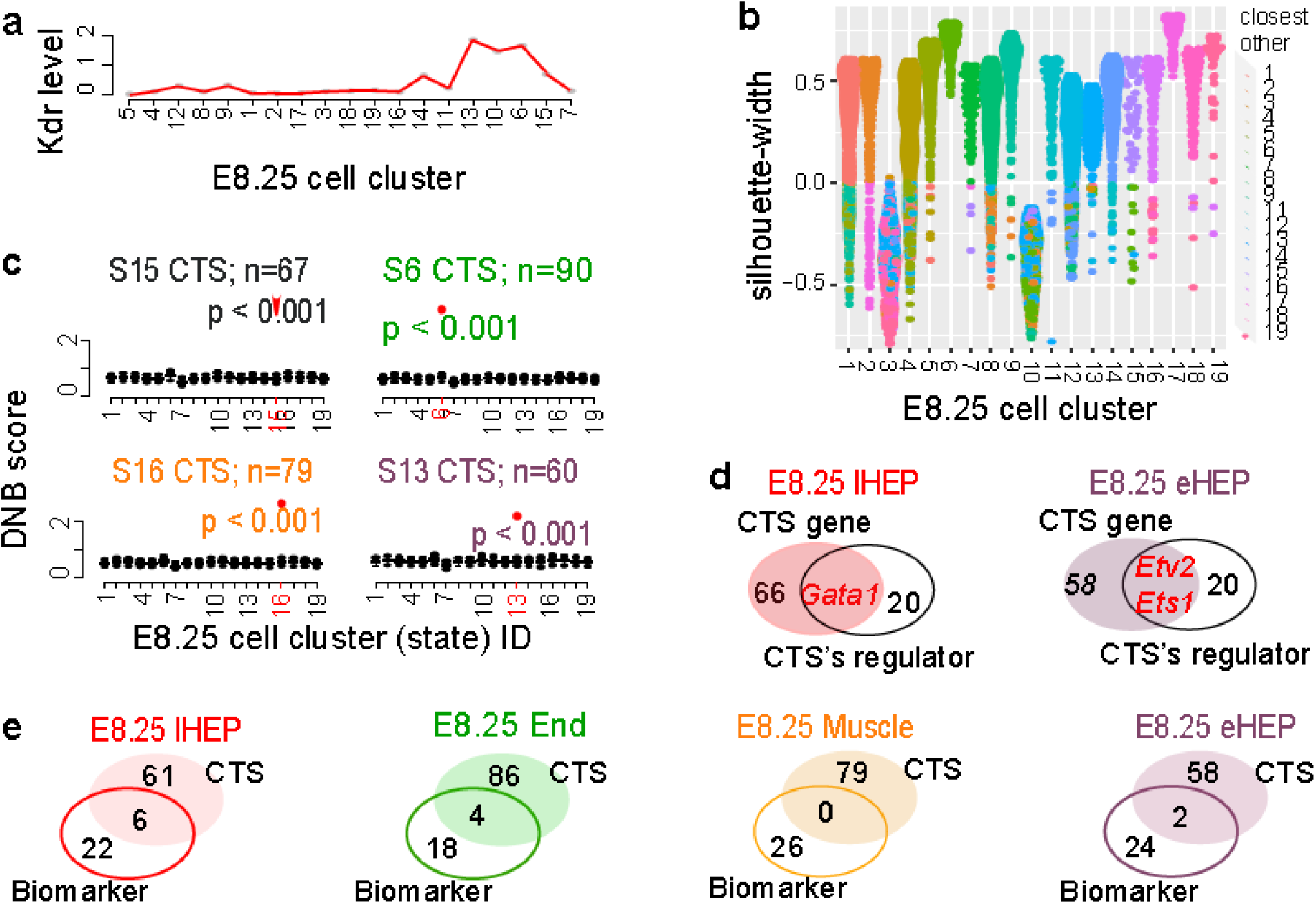
Analysis of the single-cell expression profiles of mesoderm in vivo (E8.25, 2019, GSE87038) **a**, Lines showing the average expression levels of *Kdr* in each of the 19 cell states. **b**, Distribution of the approximate silhouette width (y-axis) across 19 cell states (x-axis) of the dataset. Each point represents a cell and is colored with the identity of its own cluster if its silhouette width is positive, or that of the closest other cluster if the width is negative. **c**, Bar plots illustrate the DNB scores in 19 E8.25 states (S, GSE87038) for each identified CTS at its representative state (red dot), respectively, compared to the DNB scores of random genes (boxes, 1000 runs). **d**, Venn-diagram comparing the genes of dual roles (up-regulated biomarkers and CTS in panel a), with the up-regulators of the CTS genes. **e**, Venn-diagram for each CTS showing, up-regulated biomarkers of the representative critical transition state. lHEP: later haemato-endothelial progenitor; End: endothelial; eHEP: early HEP.

**Figure S3.**
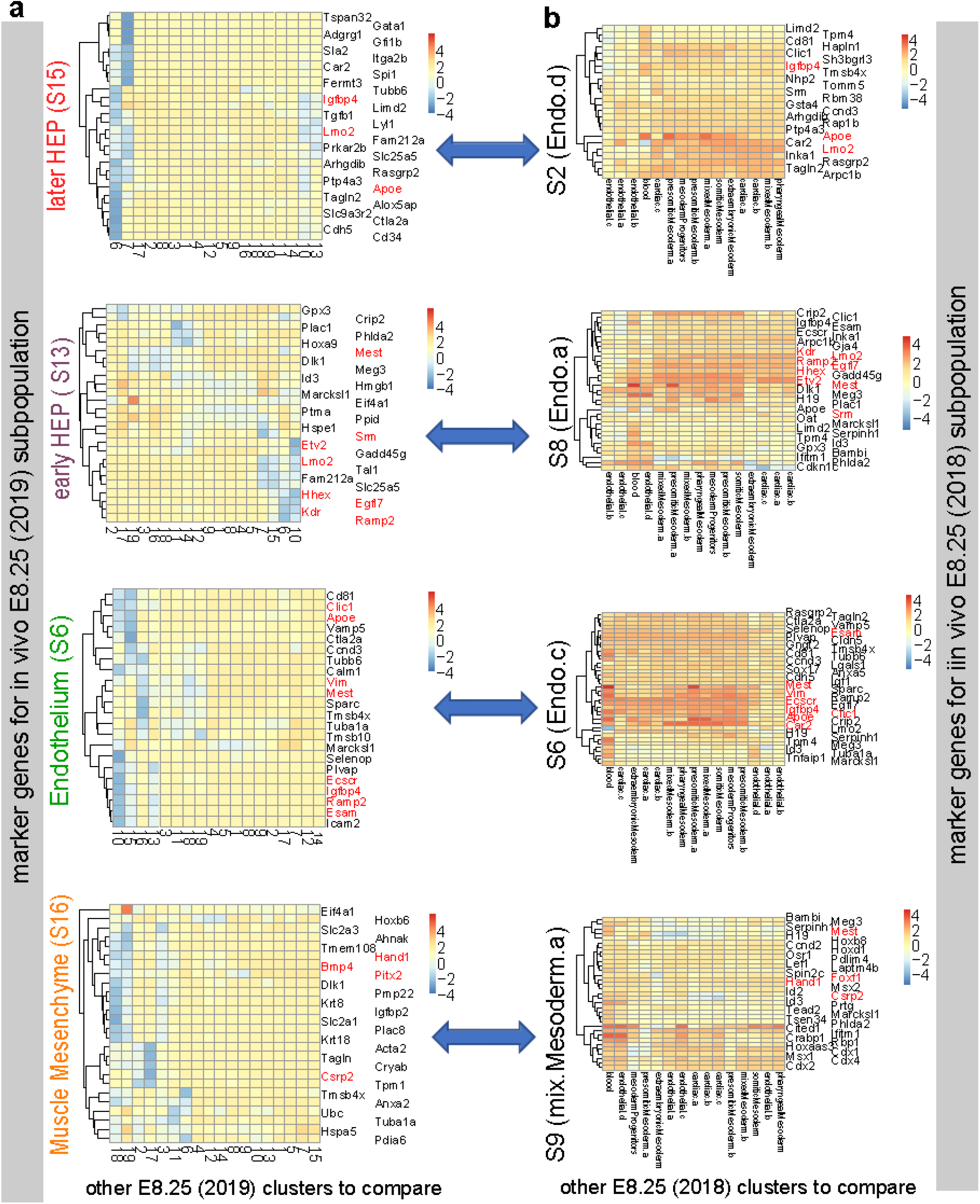
Comparing the up-regulated marker genes between two E8.25 datasets. **a**, Heatmap showing the top-5 ranked regulated genes each identified E8.25 bifurcation state over other E8.25 states (2019, GSE87038). Euclidean distance was measured, and normalized log counts were centered and scaled in the row direction. Pairwise comparisons between cell states were run using the Wilcoxon rank-sum test. **b**, Similar to panel a but for the top 10 up-regulated markers detected from the E8.25 (2018) dataset, each shown state sharing up-regulated biomarkers with one E8.25 bifurcation states (blue arrows). Pairwise comparisons between cell states were run using the t-test. The up-regulated markers were identified as a summary logFC > 1, FDR<0.01, and rank ≤ 10, using the R package scran. In both panels, the red color of gene symbols highlights shared markers.

## 12. Appendix

Other supporting Materials are available with this article online, including five S Tables and S Methods.

